# Targeted resurrection of chromosomal arm 1RS in two elite wheat lines with 1BL/1RS translocation for improved end-use quality

**DOI:** 10.1101/2021.07.29.454297

**Authors:** Ramandeep Kaur, Guriqbal Singh Dhillon, Amandeep Kaur, Sarabjit Kaur, Puneetinder Kaur, Diljot Kaur, Aman Kumar, Rohtas Singh, Gurvinder Singh Mavi, Satvir Kaur Grewal, Parveen Chhuneja, Satinder Kaur

## Abstract

1BL/1RS translocation is widely used around the world to enhance wheat yield potential, resistance to various diseases, and adaptation. However, the translocation is combined with inherent quality problems associated with reduced dough strength and dough stickiness due to the presence of *Sec-1* on proximal end and absence of *GluB3/GliB1* on distal end. Two NILs, one carrying the distal (1RS^RW^) and the other carrying the proximal (1RS^WR^) fragment from 1BS, in background of Pavon were used for transferring these two loci in yellow rust resistant version of two elite wheat varieties PBW550+*Yr5* and DBW17+*Yr5*. Foreground and background marker assisted selection was done for the *Sec-1-* and *GluB3*+ alongwith *Lr26/Yr9/Sr31*, *Pm8* and 1RS loci in the advancing generation. BC_2_F_5:6_ NILs with absence of *Secalin* and presence of *GluB3/GliB1* loci were evaluated for two years in replicated yield trial. A positive correlation of thousand grain weight (TGW), harvest index (HI), and tiller number per meter (TNpM) with yield (YD) with significant GxE effect was observed. Further multivariate analysis of these traits contributed maximum to the effective yield. Thirty promising NILs were identified with *Sec-1-/GluB3+* alongwith with high yield contributing parameters.

## Introduction

Wheat (*Triticum aestivum* L., 2n=6x=42, AABBDD), the most important staple food globally, is the major food crop next to rice in India providing 50 percent of the total calories and 60 percent of the total proteins (Grote et al. 2021). It is staple food of nearly 2.5 billion of the world population and has an estimated 137.5 million metric tons global production from 23.9 million hectares (mha) (http://www.fas.usda.gov). With the growing wheat industry and change in consumer preferences, the efforts for quality wheat production leading to improved baking and confectionery products have gained much attention (Kanojia et al. 2018).

The use of bread and bakery products can be traced back to ancient times (10,000 BC) (Valavanidis 2018). More than 9 billion kg of bread products are produced annually, with an average consumption of 41 to 303 kg per year per capita (Dong and Karboune 2021). These products are an important source of energy and a significant reservoir of protein, complex carbohydrates (mainly starch), dietary fiber, vitamins (especially B vitamins), and minerals. The bread and other cereal-based products form the base of the food pyramid and bread consumption is recommended in all dietary guidelines (Kostyuchenko et al. 2019). In the present times, the bakery is one of the fastest growing food industries, playing a significant role in economic growth, implying that wheat flour quality is of utmost importance for sustainability (Longin et al. 2020).

The quality of the wheat grain-based products is determined mainly by prolamins or storage proteins that account for 45 to 80 % of the total proteins in the grain of modern wheat cultivars. The quality of dough is the cumulative effect of total protein content, the ratio of gliadins (extensibility) to the glutenins (elasticity) (Meenakshi and Khatkar 2005; Suchy *et al.* 2003). The most widely accepted classification puts prolamins into three structural/functional classes: high molecular weight gluten subunits (HMW-GS), low molecular weight glutenin subunits (LMW-GS), and gliadins. The locus *Glu-A1, Glu-B1*, and *Glu-D1* on long arm of chromosomes 1A, 1B and 1D respectively, code for HMW-GS while *Glu-A3, Glu-B3*, and *Glu-D3* on short arm of group 1 chromosomes codes for LMW-GS (Wang et al. 2020). The *Gli-1, Gli-3, Gli-5, Gli-6* locus on the short arm of 1B chromosome code for low molecular weight gliadins with *Gli-1* being tightly linked with *Glu-B3* locus. The presence of polymeric HMW-GS and LMW-GS is positively correlated with good bread-making quality. In the gluten complex, HMW-GS and LMW-GS covalently interact with each other by inter-molecular disulfide bonds, thus exist as glutenin macro polymers (GMPs). Glutamine-rich repetitive sequences that comprise the central part of these subunits are actually responsible for the elastic properties due to extensive arrays of interchain hydrogen bonds (Anjum et al. 2007). LMW-GS contain a long repetitive domain that facilitates the formation of more α - helices and β-strands, and confers superior gluten structure and bread making quality (Wang et al. 2016). Gliadins exist mainly as monomers, and interact non-covalently with GMPs. They act as plasticizer to modify the extensibility of gluten and dough and thus the end-use traits (Wang et al. 2017). The balanced ratio of these monomeric to polymeric proteins provide the required amount of strength and viscosity for the dough (Wang et al. 2008). The loss of any of these loci will result in misproportion which will reflect in the form of weakness and stickiness of the dough (Oak and Tamhankar 2017) due to change in sedimentation value and gluten content. Higher sedimentation values indicate the higher gluten strength; hence the improved gluten quality and the gluten content indicates the quantity of gluten. The gliadins affect the loaf volume potential and dough viscosity, while glutenins affected dough development time and loaf volume (Kaur et al. 2020).

Owing to the efforts of increasing the wheat production worldwide, the introgression of rye chromosome into the wheat cultivars is quite common from long time. The 1BL.1RS translocation is one such extensively employed alien chromatin in the history of bread wheat breeding programs to improve grain yield potential (Li et al. 2020). In this translocation, the short arm of wheat 1B chromosome has been replaced by the short arm of rye 1R chromosome (1BL.1RS translocation) by chromosomal rearrangement (Lukaszewski 2014). A worldwide list of 2470 wheat cultivars and experimental lines that carry alien introgressions has been compiled by Schlegel 2014 (Crespo-Herrera et al. 2017).

The 1RS chromosome carry many important loci, resistance against leaf rust (*Lr26*), stem rust (*Sr31*) (Mago et al. 2005), stripe rust (*Yr9*), and powdery mildew (*Pm8*) (Ren et al. 2009). Moreover, the 1RS chromosome is also known to enhance the yield potential and wide range of environmental adaptability of wheat (Ren et al. 2016). It also increases above-ground biomass, deep root system, canopy water status, and abiotic stress tolerance particularly, drought tolerance (Ehdaie et al. 2003; Sharma et al. 2009; Howell et al. 2014).

Despite such a remarkable contribution of this short arm of rye, inherent quality problems associated with reduced dough strength and dough stickiness (‘sticky dough syndrome’) have been reported in wheat. This was due to presence of *Sec-1*locus on proximal part coinciding with the absence of *Glu-B3/Gli-1* loci on distal part of 1RS chromosome arm (Zhao et al. 2011). The translocation has led to the disparity in ratio of monomeric to polymeric proteins as the secalin proteins encoded by 1RS are monomeric and highly soluble in water leading to the formation of sticky dough (Barak et al. 2012). Also, due to the absence of cysteine residues in secalin protein (which were abundant in LMW-GS), they could not form comparable gluten like polymer as cysteine plays role in intermolecular and intramolecular disulphide bonding required for proper dough making. It has led inferior bread quality of low bread volume, poor mixing tolerance, and dough stickiness (Sharma et al. 2020). The deletion of LMW-GS encoding *Glu-B3* loci significantly reduced gluten strength and bread making quality (Wang et al. 2016).

The deleterious of effects 1RS chromosome on grain processing quality traits were targeted by removal and replacement of two interstitial rye segments in 1RS arm with wheat chromatin: a distal segment to introduce the *Glu-B3*locus from wheat and a proximal segment to remove the rye *Sec-1* locus. Howell et al. (2014) successfully generated two NILs in background of cultivar Pavon, one with absence of *Sec-1* locus (called 1RS^WR^) and other with presence of *Glu-B3* locus (called 1RS^RW^) in 1RS chromosomal arm. In the present study, these two NILs were crossed and backcrossed with two important 1RS.1BL chromosome carrying wheat cultivars PBW550 and DBW17, to generate improved lines with absence of *Sec-1* locus and presence of *Glu-B3* locus through marker-assisted backcross breeding (MABB).

## Material and methods

### Plant material

Plant material includes two donor NILs in Pavon background with 1RS.1BL translocation *viz.*, Pavon 40:9 carrying *Glu-B3/Gli-B1* locus of wheat at the distal end of 1RS arm and Pavon 44:38 carrying wheat segment in place of *Sec-1* locus of rye at proximal end of 1RS arm, kindly provided by Prof. Dubcovsky from UC Davis, California. Pavon 40:9 NIL was *Glu-B3+/Sec-1+*, while Pavon 44:38 was *Glu-B3-/Sec-1*-(Fig. S1). Stripe rust resistant versions (owing to introgression of gene *Yr5* on long arm of chromosome 2B) of elite wheat cultivars PBW550 and DBW17 having 1BL/1RS translocation and were used as the recurrent parent. PBW550 (WH594/RAJ3856//W485) has been developed and released by PAU for cultivation under timely sown irrigated (TSI) conditions of North western Plain Zone (NWPZ) of India. DBW17 (CMH79A 95/3*CNO79// RAJ3777) has been developed by Indian Institute of Wheat & Barley Research (IIWBR), Karnal and has been released for cultivation under TSI in NWPZ.

### Introgression for wheat loci *Glu-B3+* and *Sec-1-* in PBW550 and DBW17

Pavon 40:9 and Pavon 44:38 were crossed as male with PBW 550 and DBW 17 to develop *Glu-B3+* and *Sec-1-* version of these two widely grown, popular wheat varieties (Fig. 1). F_1_ was backcrossed twice, and BC_2_F_1_ thus generated were selfed to obtain BC_2_F_2_, BC_2_F_3_, BC_2_F_4_, BC_2_F_5_, BC_2_F_6_ generations. Progenies were advanced by shuttle breeding between Punjab Agricultural University (PAU), Ludhiana (30.91°N, 75.85°E), between November-May (called main season - MS) and at Regional Research Stationof PAU at Keylong, Himachal Pradesh (32.71°N, 77.32°E) between May-October (called offseason OS).

**Fig.1:**
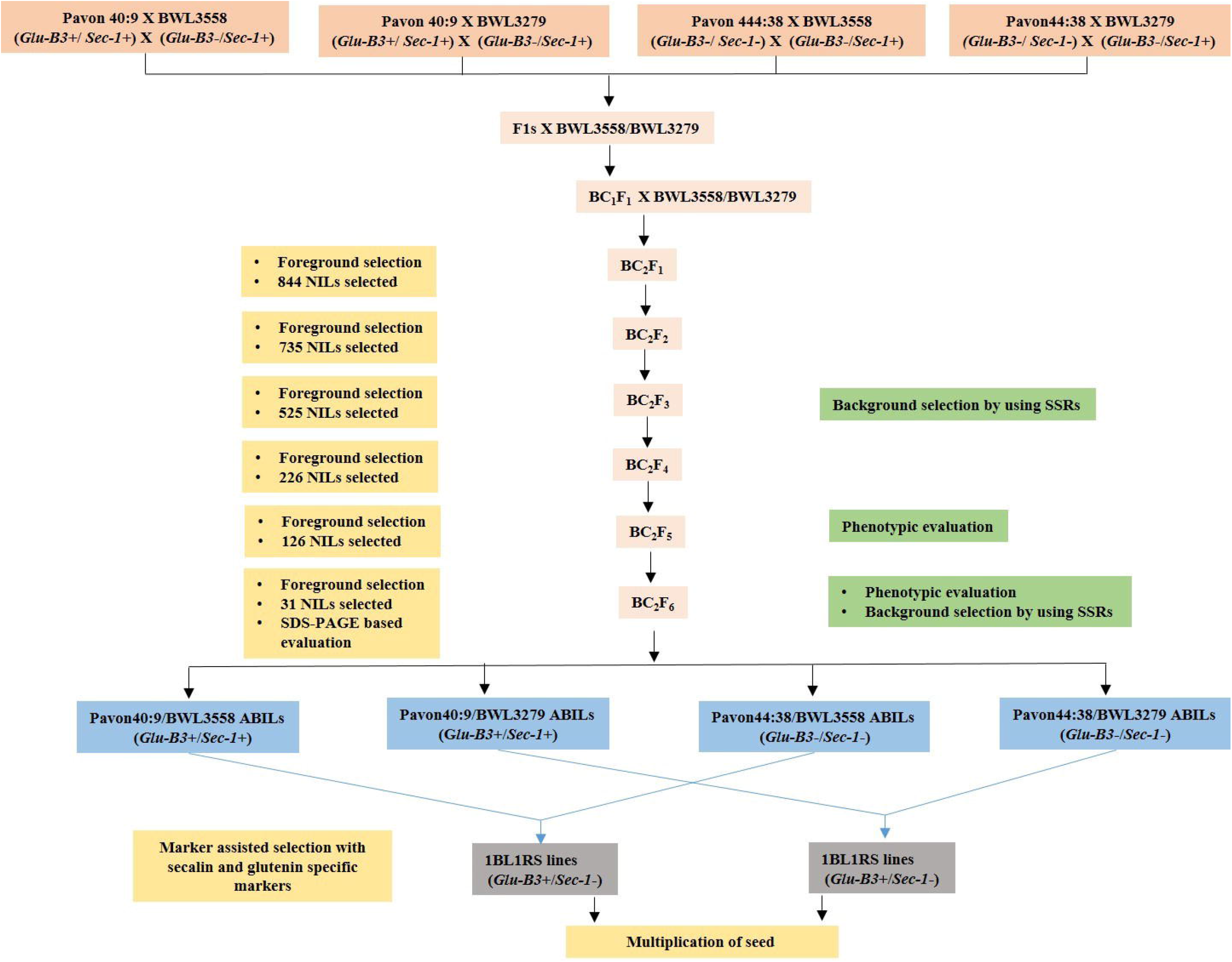
Breeding scheme of the recombinant 1BL1RS wheat lines

### Marker-assisted foreground selection

The detail of the markers used for foreground selection is given in Table S2. Two different markers, *omega p3-p4* (amplifying the secalin gene of rye) and *wpt1911* (amplifying its wheat counterpart), present on the proximal part of the short arm of 1RS and 1BS chromosome respectively were used to confirm the absence of the *Sec-1* gene and presence of wheat chromatin. *Glu-B3/Gli-B1* gene was selected with the help of *markerPsp3000.* Stripe rust resistance gene of recurrent parents, *Yr5*, was followed using the *STS-7/8* marker. Genomic DNA isolation from parental wheat genotypes and the back cross progenies was done from the young leaf tissue using the CTAB method with some modifications (Table S1) (Saghai Maroof *et al.* 1984). DNA was quantified on 1% agarose gel. The Polymerase chain reaction (PCR) was carried out in 20μl reaction volume containing 35–50 ng of genomic DNA, 1-unit Taq polymerase (Homemade), 0.15 mM of each dNTPs,1.5mM of MgCl_2_, 0.38 μM of forward and reverse primers, and 1X PCR buffer (10 mM Tris–HCl pH 8.4). The PCR products were resolved using 2% agarose gel.

### Background selection

For background recovery of the respective recurrent parent, 20-30 SSR markers were selected from each of the B-genome chromosomes of wheat. Since the present study is focused on the recovery of chromosome 1RS/1BL, a set of 50 SSR markers were selected from 1BL to ensure its recovery. For background recovery of chromosomal arm 1RS, (except two targeted loci) marker-assisted selection of the cluster of linked genes *Lr26/Yr9/Sr31* (with *Iag95* marker), *Pm8* (using *Sfr43* marker), and 1RS (using rye F3/R3marker) were done (Table S2).

### Validation through SDS-PAGE analysis

Seed storage proteins were sequentially extracted according to Smith and Payne (1984) with some modifications. Briefly, first albumins and globulins were removed from flour for the extraction of gliadin and glutenin using 1.5 M DMF (dimethylformamide) glutenin extraction buffer {50% Isopropanol, 50mM Tris-HCl (pH 7.5), 1% dithiothreitol (DTT)}. The presence/absence of *Glu-B3* encoded low molecular weight glutenin subunit (LMW-GS) in the range of 42-50kDa proteins and *Sec-1* encoded secalin protein in the range of 42-55kDa were detected through SDS-PAGE (Walker 1996). The electrophoresis unit was run for 3 hours for glutenin samples and 10 hours for gliadin samples at 50mA. After the electrophoresis, the gel was stained for 2 hours with staining solution and for de-staining, the gel was kept in a de-staining solution at shaker with minimal speed for 2 to 3 hours. The absence and presence of secalin and glutenins respectively, were observed with respect to the Pavon40:9 and Pavon44:38.

### Evaluation for agro-morphological traits

BC_2_F_5_ and BC_2_F_6_ derived NILs from the two crosses along with the two donor and two recurrent parents were evaluated in in alpha lattice design sown in three replications, during 2018-19 MS and 2019-20 MS, respectively, at PAU, Ludhiana. Each entry in trial was sown in four rows of 1.5m length, with row-to-row spacing of 25cm, and plant to plant spacing of 10cm. Agro-morphological traits of plant height (PH in cm), spikelets per spike (SS), spike length (SL) (cm), tillers per meter (TNpM), 1000 grain weight (TGW) (g), yield per plot (yield per plot YD) (g), harvest index (HI) (%) were recorded.

### Screening for stripe rust

Yellow rust severity was also recorded for both the years. Uredinospores of known stripe rust (100S119, 78S84) races mixed with local inoculum collected from farmer’s fields were sprayed to create the artificial rust epidemic. The disease severity was scored as the percentage of leaf area covered by rust following the modified Cobb’s scale, as developed by Peterson et al. (1948) as no infection (0), resistant (R), moderately resistant (MR), moderately susceptible (MS) and susceptible (S).

### Statistical analysis

Descriptive analysis and variability studies were done using Summary Toolsv0.9.4 package in R-studio (Comtois 2020). The calculation of adjusted mean values (BLUPs) was done using META-R version 6.0 (Alvarado et al. 2016). Comparison of adjusted mean values (BLUPs) was made between genotypes to the respective recurrent parent separately for both the environments. The genotypic coefficient of variability (GCV) and the phenotypic coefficient of variability (PCV) were estimated according to the the Burton and Devane (1953) by using formula

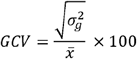

where 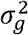 is the error/residual variance, and x is the mean of all genotypes. The phenotypic coefficient of variability (PCV) was calculated using the equation

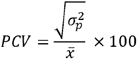

where 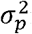 is the phenotypic variance. The environmental coefficient of variability (ECV) was calculated using the equation

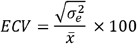

where 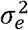 is the genotypic variance. The broad-sense heritability estimated the quality of the breeding program for the traits and the environments. The broad-sense heritability and genetic advance over mean were estimated using the formula by Allard (1960).

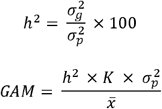

Where K is selection differential at 5% intensity of selection (K=2.06). The correlations were calculated as simple pairwise Pearson’s correlations among traits. It is defined as:

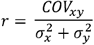

Where *COV_xy_* is the covariance between trait x and trait y, 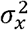 is the variance of trait x, and 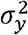 is the variance of trait y. The coefficient of skewness (β_1_) and kurtosis (β_2_) is as:

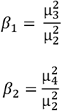

Where 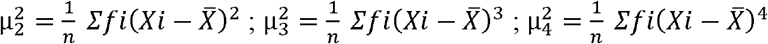

The relationship among the traits was studied by principal component analysis using FactoMineR v2.4 (Lê et al. 2008) and FactoExtra v1.0.7 (Kassambara and Mundt 2020) in Rstudiov4.0.3. The principal components were plotted for the two years as biplots of eigenvectors. Structural equation modeling (SEM) was done using the package lavaan v 0.6-7 (Rosseel, 2012) and visualized using package semPlot v1.1.2 (Epskamp et al., 2019) to identify the direct and indirect contributors of yield.

## Results

### Generation and validation of genetic material

Two crosses s were initiated in year 2015, by crossing recurrent parent PBW550/DBW17 X Pavon 44:38/Pavon 40:9 for generation of two type of stable NILs, one with absence of secalin (*Sec-1-*) gene and other with presence of *Glu-B3*in 1RS chromosomal arm of two recurrent parents. In the advancing generations, plants were selected for genes *Sec-1-*, 1RS, *Pm8* and *Lr26/Yr9/Sr31* in cross Pavon 44:38X PBW550/DBW17 and for *GluB3*, 1RS, *Pm8* and *Lr26/Yr9/Sr31* in cross Pavon 40:9 X PBW550/DBW17. Number of plants selected in advancing backcross generations in both the crosses are given in Table S1 and schematic representation for the development of improved 1BL/1RS is represented in Fig.1. In 2016 OS, foreground selection was done on 1026 BC_1_F_1_plants and for absence of *Sec-1-* gene and for presence of *Glu-B3+* along with three genes of chromosomal arm 1RS. The selected plants were backcrossed with respective recurrent parents and 1925 BC_2_F_1_ plants were sown in 2016-17 MS, again MAS was done for four genes in homozygous/heterozygous form in both the crosses. Background selection was done on the plants positive for four genes (homozygous/heterozygous) and a recurrent parent recovery of 75% or more was selected. In 2017 OS, BC_2_F_2_ generation was sown as plant to row progenies and two plants/progeny, homozygous for atleast three of four targeted genes were selected. In 2017-18 MS, BC_2_F_3_ was again sown as plant to row and progenies homozygous for 3-4 of targeted genes were selected and three single plants harvested from selected progenies. In 2018 OS, BC_2_F_4_ sown in plant to row and again selection was made for four genes in homozygous form, selecting three plants/progeny. Replicated yield trial of the developed BC_2_F_5_ and BC_2_F_6_ NILs, in both of crosses was done in 2018-19 MS and 2019-20 MS respectively and here again MAS was done for four targeted genes selecting five plants/progeny of the selected progenies. 45 best performing BC_2_F_6_ NILs were selected and validated for the presence of glutenin/gliadin (42-50kDa) LMW proteins encoded by *Glu-B3* gene and absence secalin protein (42-55kDA) on SDS base PAGE (Fig 2). 30 NILs were finally selected based on the yield related traits having 1RS chromosomal arm substituted with loci *Sec-1* and *Glu-B3* from wheat 1BS chromosomal arm.

**Fig. 2:**
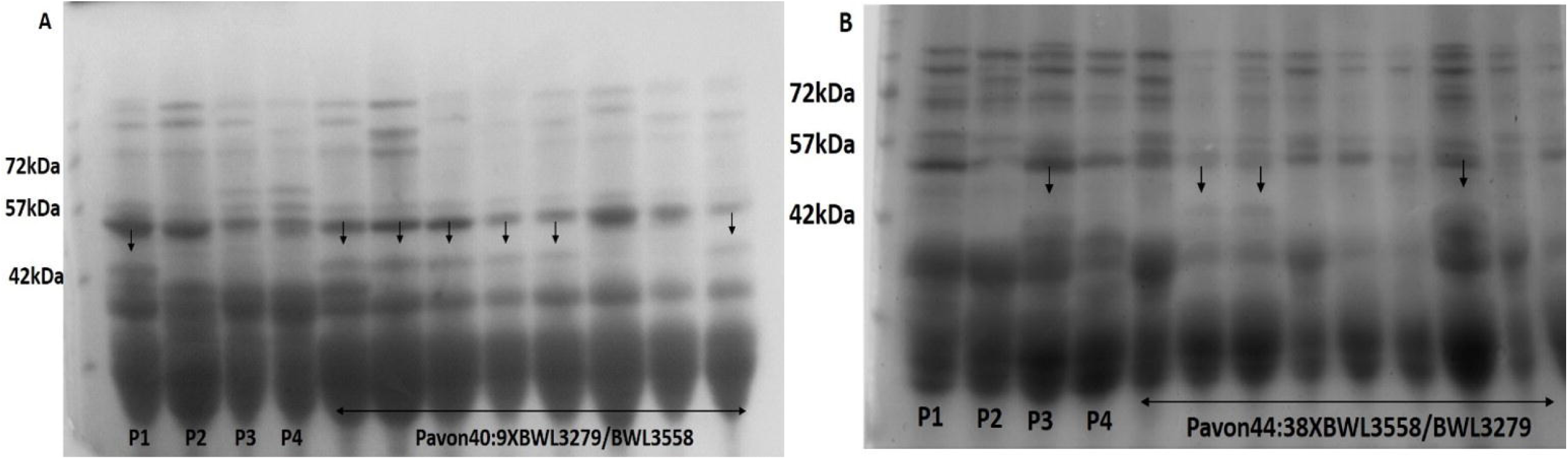
Validation of presence of A) Glu-B3 and B) Sec-1 locus in Pavon 40:9XBWL3558/BWL3279 and Pavon 44:38XBWL358/BWL3279 BC_2_F_6_ near isogenic lines (NILs) developed through marker-assisted introgression. Fig 1A: P1-Pavon40:9, P2-Pavon44:38, P3-BWL3558, P4-BWL3279; Fig 1B: P1-PBW343, P2-Chinese spring, P3-BWL3558, P4-Pavon44:38

### Phenotypic evaluation

The phenotypic evaluations for yield was done on BC_2_F_5_ and BC_2_F_6_ NILs for two consecutive years from 2018-20 in alpha lattice design. In 2018-19, 172 BC_2_F_5_ NILs in the background of PBW550 (103) and DBW17 (69) without *Sec-1* gene, 54 NILs with *Glu-B3*+ gene in the background of PBW550 (39) and DBW17 (15) were evaluated. 126 selected BC_2_F_6_ NILs were evaluated in the year 2019-20 with 110 NILs without *Sec-1* gene (in background of PBW550 - 61 and DBW17– 49) and 16 NILs with *Glu-B3* gene (in background of PBW550 - 11 and DBW17 – 5) (Table S1).

The adjusted values based on ANOVA of each trial were compared with respective recurrent parents for selections. Significant variation was observed within the NILs, and the least variation was observed across the years (Fig. 3). The yield-related traits, *i.e.*, TGW, TNpM, HI, and yield, showed significant improvements signifying the positive effect of selection across the two years (Fig. 4).

**Fig 3:**
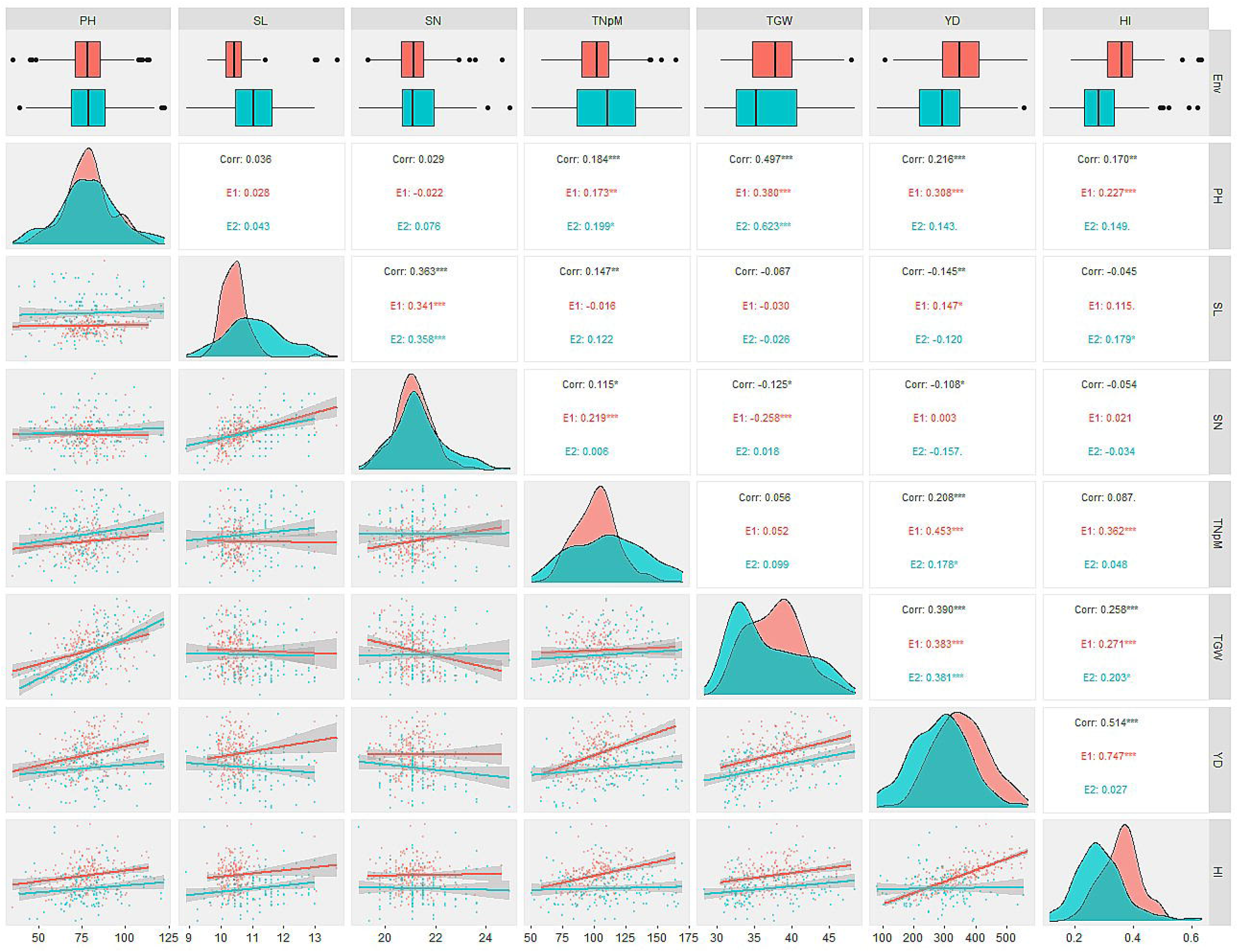
Phenotypic evaluation of near isogenic lines (NILs) generated in the present study across two generations BC_2_F_5_ and BC_2_F_6_

**Fig. 4:**
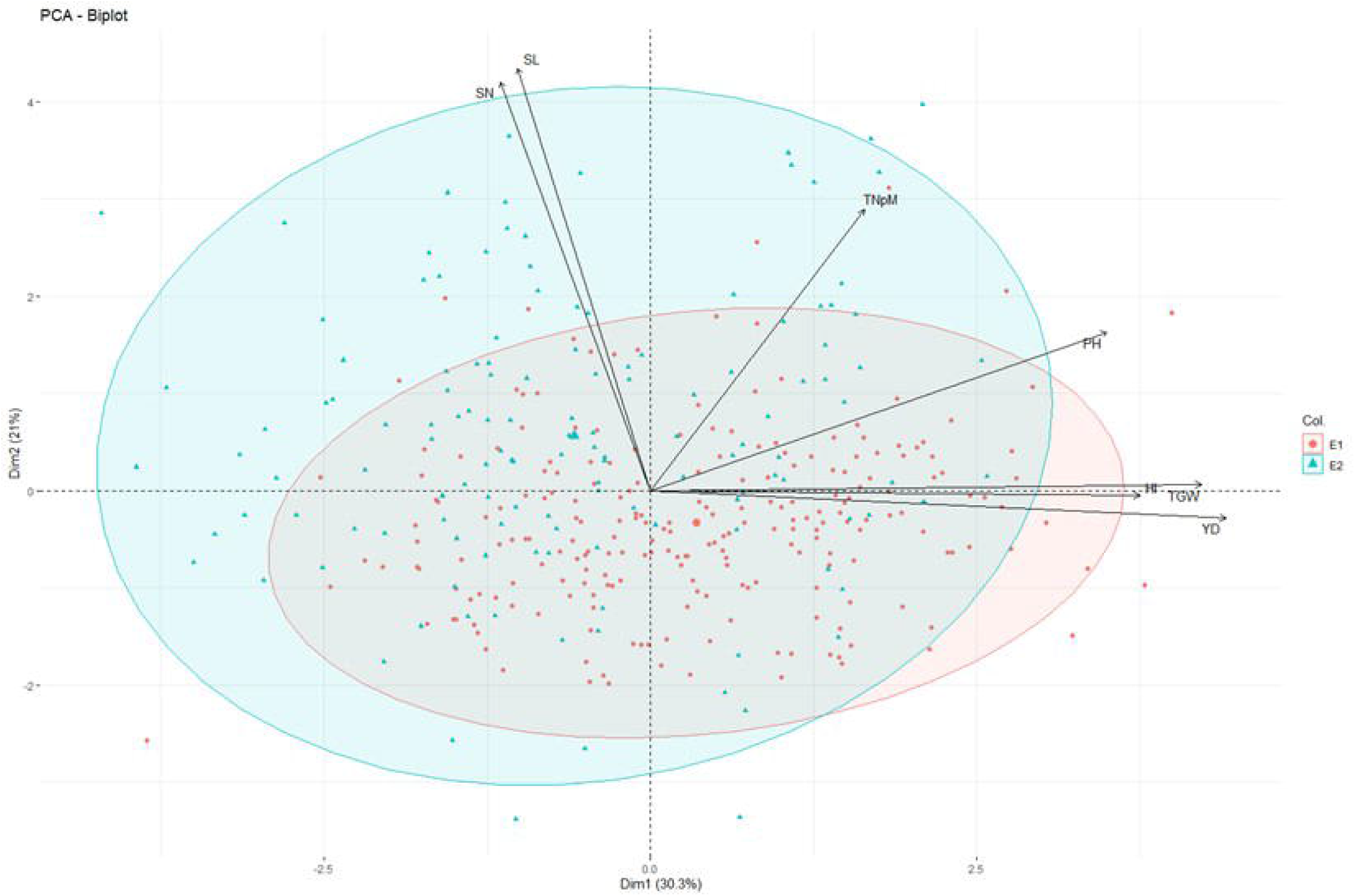
Principal component analysis of near isogenic lines (NILs) generated in the present study across two generations BC_2_F_5_ and BC_2_F_6_

The average range of PH was from 35.37-113.66cm and 39.13-122.18cmcompared to the recurrent parents PBW550 (84.18cm; 86.92cm) and DBW17 (77.36cm; 80.99cm) during 2018-19 and 2019-20 season, respectively. SL ranged from 9.57-13.68cm and 8.91-12.99cm with recurrent parents PBW550 (10.24cm; 11.24cm) and DBW17 (9.64cm; 11.83cm; during 2018-19 and 2019-20 season, respectively).For the recurrent parents, PBW550 and DBW17, SN was 20.35; 20.25 and 21.38; 21.10 with the range of NILs from 19.34-24.64 and 18.97-24.94 during 2018-19 and 2019-20 season, respectively. TNpM ranged from 58.31-164.65 and 50.79-169.87compared with the recurrent parentsPBW550 and DBW17being 96.49; 105.84 and 106.20; 114.58 during 2018-19 and 2019-20 season. TGW for NILs ranged from 30.53-48.02g and 28.38-48.51g, with the recurrent parents PBW550 and DBW17 TGW being 43.00g; 40.16g and 39.77g; 37.86g during 2018-19 and 2019-20 season respectively. Similarly, YD ranged from 108.38-568.60g and 83.67-554.56g during 2018-19 and 2019-20 season with recurrent parentsPBW550 and DBW17 YD of 409.80g; 402.25g and 419.96g; 347.45g for both the seasons 2018-19 and 2019-20. HI ranged from 0.19-0.63% and 0.11-0.62% compared with the recurrent parents PBW550 and DBW17 HI being 0.40%; 0.47% and 0.41%; 0.38% during 2018-19 and 2019-20 season respectively (Table 1).

**Table 1.** Phenotypic variabilities of near isogenic lines (NILs) generated in the present study across two generations BC_2_F_5_ and BC_2_F_6_

| **Trait** | **Env** | **BWL3558** | **BWL3279** | **Pavon40:9** | **Pavon44.38** | **Range** | **Mean** | **StdDev** | **Skewness** | **Kurtosis** |
| --- | --- | --- | --- | --- | --- | --- | --- | --- | --- | --- |
| PH | E1 | 84.18 | 77.36 | 66.25 | 79.91 | 35.37-113.66 | 78.71 | 13.69 | 0.03 | 0.29 |
|  | E2 | 86.92 | 80.99 | 72.09 | 81.98 | 39.13-122.18 | 78.97 | 17.13 | 0.15 | 0.03 |
| SL | E1 | 10.24 | 9.64 | 10.15 | 10.00 | 9.57-13.68 | 10.45 | 0.48 | 2.68 | 14.20 |
|  | E2 | 11.24 | 11.83 | 11.63 | 12.80 | 8.91-12.99 | 11.07 | 0.90 | 0.16 | -0.42 |
| SN | E1 | 20.35 | 21.38 | 19.83 | 20.64 | 19.34-24.64 | 21.14 | 0.75 | 0.57 | 1.91 |
|  | E2 | 20.25 | 21.10 | 19.40 | 18.12 | 18.97-24.94 | 21.35 | 1.05 | 0.65 | 0.61 |
| TNpM | E1 | 96.49 | 106.20 | 107.82 | 87.75 | 58.31-164.65 | 101.72 | 17.07 | 0.38 | 0.56 |
|  | E2 | 105.84 | 114.58 | 112.32 | 104.54 | 50.79-169.87 | 110.86 | 28.61 | 0.08 | -0.79 |
| TGW | E1 | 43.00 | 39.77 | 31.98 | 39.82 | 30.53-48.02 | 37.72 | 3.63 | 0.24 | -0.41 |
|  | E2 | 40.16 | 37.86 | 37.53 | 38.04 | 28.38-48.51 | 36.84 | 4.78 | 0.56 | -0.92 |
| YD | E1 | 409.80 | 419.96 | 410.73 | 360.56 | 108.38-568.60 | 353.62 | 86.40 | 0.11 | -0.30 |
|  | E2 | 402.25 | 347.45 | 421.60 | 441.05 | 83.67-554.56 | 283.91 | 92.27 | 0.11 | -0.03 |
| HI | E1 | 0.40 | 0.41 | 0.41 | 0.30 | 0.19-0.63 | 0.36 | 0.07 | 0.41 | 1.10 |
|  | E2 | 0.47 | 0.38 | 0.40 | 0.32 | 0.11-0.62 | 0.29 | 0.09 | 0.93 | 1.88 |

Plant height (PH), Spike length (SL), spikelet no. per spike (SN), tiller number per meter (TNpM), thousand grain weight (TGW), yield per plot (YD), and harvest index (HI).

All the traits exhibited positive skewness for both years, suggesting that the genetic gain obtained is through intense selection (Table 1). PH exhibited skewness of 0.03 and 0.15 with leptokurtic distribution (0.29, 0.03) for both years, with a decrease in value for the second year. SL showed positive skewness of 2.68 and 0.16 with platykurtic distribution (−0.4.2) in the second year compared to the first year (14.20). SN showed positive skewness of 0.57 and 0.65 with leptokurtic distribution for both years (1.91, 0.61). The TNpM exhibited positive skewness of 0.38 and 0.08 with leptokurtic distribution (0.56) for the first year and platykurtic distribution for the second year (−0.79) for the second year. The TGW showed positively skewed values 0.24 and 0.56 with platykurtic distribution for both years (−0.41, −0.92). While, the yield showed positively skewed values 0.11 and 0.11 with platykurtic distribution for both the years (-0.30, −0.03).The HI showed positive skewness of 0.41 and 0.93 with leptokurtic distribution for both years (1.10, 1.88).

### Genetic variability, heritability, and genetic advance

The genetic variability parameters GCV and PCV are characterized to be low from 0-10%, moderate from 10-20%, and high above 20%. The estimates of variability are given in Table 2. In the study, moderate to high GCV and PCV were observed for all the traits except SL and SN. The GCV and PCV ranged from 17.86-25.55% and 19.62-26.00% for PH, 6.78-10.82% and 14.52-19.18% for SL, 4.58-6.45% and 7.80-1059% for SN, 16.99-26.92% and 17.74-28.08% for TNpM, 10.01-13.22% and 11.14-14.48% for TGW, 24.65-32.68% and 25.16-33.64% for yield, 23.28-47.35% and 32.65-55.90% for HI. The high coefficient of variation values suggests high variability among the population. The GCV value for all the traits is less than the ECV value. The more the value of GCV than ECV, the less is the environment effect.

**Table 2.** Genotypic variabilities of near isogenic lines (NILs) generated in the present study across two generations BC_2_F_5_ and BC_2_F_6_

| Trait | Env | GCV | ECV | PCV | H<sup>2</sup> | GAM | LSD | CV |
| --- | --- | --- | --- | --- | --- | --- | --- | --- |
| PH | E1 | 17.86 | 08.11 | 19.62 | 91.03 | 36.79 | 9.90 | 8.11 |
|  | E2 | 25.56 | 04.74 | 26.00 | 98.31 | 52.65 | 6.05 | 4.74 |
| SL | E1 | 06.78 | 12.84 | 14.52 | 46.69 | 13.96 | 1.45 | 12.84 |
|  | E2 | 10.82 | 15.84 | 19.18 | 56.41 | 22.29 | 2.16 | 15.84 |
| SN | E1 | 04.58 | 06.32 | 07.80 | 58.72 | 09.44 | 1.68 | 6.32 |
|  | E2 | 06.45 | 08.40 | 10.59 | 60.91 | 13.29 | 2.30 | 8.40 |
| TNpM | E1 | 16.99 | 05.11 | 17.74 | 95.77 | 35.00 | 8.22 | 5.11 |
|  | E2 | 26.92 | 07.98 | 28.08 | 95.87 | 55.45 | 14.12 | 7.98 |
| TGW | E1 | 10.01 | 04.89 | 11.14 | 89.86 | 20.63 | 2.85 | 4.89 |
|  | E2 | 13.22 | 05.90 | 14.48 | 91.30 | 27.23 | 3.39 | 5.90 |
| YD | E1 | 24.65 | 05.07 | 25.16 | 97.97 | 50.78 | 28.55 | 5.07 |
|  | E2 | 32.68 | 07.99 | 33.64 | 97.15 | 67.33 | 36.62 | 7.99 |
| HI | E1 | 23.28 | 22.90 | 32.65 | 71.30 | 47.96 | 0.11 | 22.90 |
|  | E2 | 47.35 | 29.71 | 55.90 | 84.70 | 97.53 | 0.17 | 35.77 |

3 Heritability broad sense( $h^2$ ), Genotypic Coefficient of Variance (GCV), Phenotypic Coefficient of 4 Variance (PCV), Residual/Environmental Coefficient of Variance (ECV) and Coefficient of 5 variation significant at $\alpha < 0.05$ (CV)

Broad sense heritability ranged from moderate to high among the characters from 46.69-98.31% across two years. Moderate to high heritability was observed for SL (46.69-56.41%) and SN (58.72-60.91%). High heritability was observed for PH (91.03-98.31%), TNpM (95.77-95.87%), TGW (89.86-91.30%), YD (97.97-97.15%) and HI (71.30-84.70%). Genetic advance is measured as low (for characters showing a GAM value of <10%), moderate (for characters showing a GAM value of 10-20%), and high (for characters showing a GAM value of >20%) (Johnson *et al.*, 1955). Moderate to high GAM values were observed for SL (13.96-22.29%) and SN (09.44-13.29%). High GAM value was observed for PH (36.79-52.65%), TNpM (35.00-55.45%), TGW (20.63-27.23%), YD (50.78-67.33%) and HI (47.96-97.53%). Although there is not much difference between the mean values of the traits for both years (Table 3), the GAM of the second year is significantly more than the first year, hence the superior selection of progenies.

**Table 3:** Selected high performing 1BLIRS near isogenic lines (NILs) for various yield component traits

| **Recurrent parent** | **PH** | **SL** | **SN** | **TNpM** | **TGW** | **YD** | **HI** |
| --- | --- | --- | --- | --- | --- | --- | --- |
| BWL3558 (PBW550+Yr5) | 86.92 | 11.24 | 20.25 | 105.84 | 40.16 | 402.25 | 0.47 |
| BWL3279 (DBW17+Yr5) | 80.99 | 11.83 | 21.10 | 114.58 | 37.86 | 347.45 | 0.38 |
| **Pavon44:38XBWL3558** |  |  |  |  |  |  |  |
| ABIL-11 | 76.37 | 12.02 | 21.96 | 142.11 | 34.53 | 365.89 | 0.40 |
| ABIL-13 | 80.24 | 12.60 | 22.81 | 70.86 | 37.34 | 251.37 | 0.59 |
| ABIL-14 | 89.23 | 12.02 | 23.66 | 158.30 | 43.21 | 390.94 | 0.49 |
| ABIL-26 | 94.49 | 10.65 | 21.10 | 105.83 | 45.12 | 369.42 | 0.50 |
| ABIL-72 | 95.82 | 10.46 | 21.10 | 92.77 | 46.87 | 433.45 | 0.23 |
| ABIL-79 | 84.61 | 10.66 | 21.10 | 113.29 | 41.01 | 382.77 | 0.32 |
| ABIL-86 | 110.6 | 12.41 | 22.38 | 133.04 | 41.93 | 456.09 | 0.34 |
| ABIL-88 | 79.00 | 10.46 | 21.52 | 124.62 | 31.10 | 367.26 | 0.37 |
| ABIL-90 | 94.50 | 10.66 | 21.10 | 105.84 | 45.12 | 369.42 | 0.50 |
| ABIL-97 | 83.95 | 11.05 | 20.68 | 114.91 | 37.99 | 439.75 | 0.33 |
| ABIL-102 | 82.31 | 10.08 | 21.10 | 83.49 | 45.28 | 537.67 | 0.35 |
| **Pavon44:38XBWL3279** |  |  |  |  |  |  |  |
| ABIL-41 | 83.62 | 11.44 | 21.96 | 147.29 | 39.47 | 360.18 | 0.32 |
| ABIL-107 | 82.42 | 11.63 | 21.10 | 133.69 | 34.67 | 384.93 | 0.32 |
| ABIL-108 | 83.04 | 10.08 | 20.25 | 100.23 | 33.85 | 360.58 | 0.30 |
| ABIL-110 | 97.79 | 9.69 | 20.68 | 144.05 | 39.16 | 448.66 | 0.36 |
| ABIL-115 | 74.73 | 11.05 | 20.68 | 108.75 | 38.91 | 409.98 | 0.35 |
| ABIL-119 | 107.9 | 11.24 | 21.53 | 138.55 | 36.58 | 366.66 | 0.34 |
| ABIL-122 | 94.17 | 10.27 | 20.68 | 126.57 | 45.10 | 424.76 | 0.30 |
| ABIL-123 | 83.95 | 10.69 | 19.40 | 134.01 | 42.67 | 381.05 | 0.36 |
| ABIL-128 | 74.73 | 11.05 | 19.83 | 109.40 | 36.47 | 395.20 | 0.32 |
| ABIL-131 | 82.86 | 11.63 | 21.53 | 125.92 | 34.81 | 360.24 | 0.37 |
| **Pavon40:9XBWL3558** |  |  |  |  |  |  |  |
| ABIL-53 | 71.10 | 10.46 | 21.10 | 90.29 | 32.65 | 321.56 | 0.27 |
| ABIL-55 | 74.07 | 10.46 | 21.10 | 105.19 | 32.54 | 328.78 | 0.24 |
| ABIL-56 | 81.98 | 10.85 | 21.10 | 137.58 | 46.23 | 304.26 | 0.25 |
| ABIL-64 | 73.17 | 12.80 | 19.83 | 136.28 | 30.97 | 300.83 | 0.26 |
| ABIL-65 | 76.04 | 10.08 | 21.10 | 85.76 | 33.36 | 295.73 | 0.24 |
| ABIL-67 | 72.42 | 11.05 | 21.10 | 127.54 | 38.53 | 311.87 | 0.36 |
| **Pavon40:9XBWL3279** |  |  |  |  |  |  |  |
| ABIL-57 | 74.73 | 11.05 | 21.10 | 94.50 | 38.34 | 264.33 | 0.35 |
| ABIL-58 | 72.76 | 10.91 | 20.25 | 73.78 | 32.19 | 367.64 | 0.37 |
| ABIL-61 | 88.90 | 9.88 | 21.53 | 107.46 | 45.27 | 414.44 | 0.27 |

### Correlation analysis

Correlation studies between various agro-morphological traits were carried out to establish their relationship with yield. For progenies, PH showed a positive correlation with yield for 2018-19 (0.308) and 2019-20 (0.143). SL showed a positive correlation with yield for 2018-19 (0.147) and a negative correlation in the year 2019-20 (−0.120). SN showed a positive correlation with yield for 2018-19 (0.003) and a negative in the year 2019-20 (−0.157). However, both TNpM and TGW showed a highly significant positive correlation with yield. TNpM showed a positive correlation of 0.453 and 0.178 for the years2018-19 and 2019-20, respectively. Similarly, TGW showed a positive correlation of 0.383 and 0.381 for the years2018-19 and 2019-20. HI showed a positive correlation with yield for both the years 2018-19 (0.747) and 2019-20 (0.027) (Fig. 3).

### Principle component analysis

Principal component analysis as an exploratory tool for data analysis. It offers details about traits by elucidating the population’s maximum variability in the given environments (Fig. 4). The eigen vectors in the first two principal components explained only 51.3% of the total variability across the environments. The high GxE effect of the lines might have occurred due to the selection of superior genotypes across the years, the difference in sowing dates, or change into year-to-year weather. Overall, the PCA showed that YD was dependent more on TGW, HI and least dependent on SL and SN. However, SN was more dependent on SL. YD was also dependent upon PH and TNpM, as discussed in correlation analysis. TGW was dependent on HI more than that of PH.

### Multivariate analysis by Structural equation modelling

The PCA showed that YD was dependent on all the characters studied during the present experiment to different extents, structural equation modelling was used to study the direct and indirect variables which determined the YD. The SEM showed that the TNpM, TGW and HI were the main direct contributors of YD (Fig. 5). The PH also showed indirect contribution towards YD through TGW and TNpM which were contributing directly towards the YD.

**Fig. 5:**
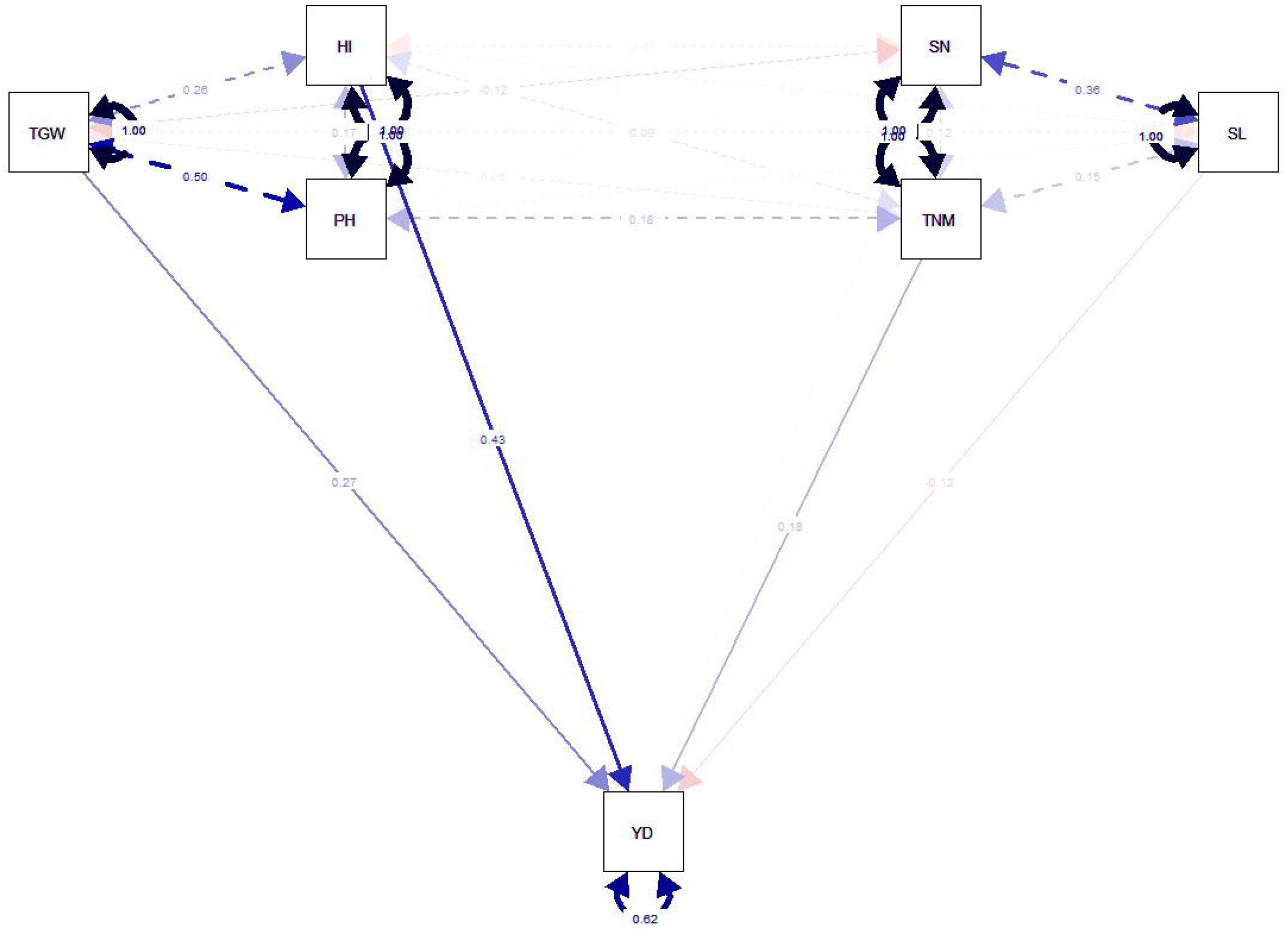
Multivariate analysis by Structural equation modeling of near isogenic lines (NILs) generated in the present study across two generations BC_2_F_5_ and BC_2_F_6_

### Identification of superior NILs

A total of 21 NILs with *Sec-1-gene* transferred by the donor parent Pavon44:38 in the background of PBW550 and DBW17showed superior performance in yield-related traits than their respective recurrent parent (Table 3). Out of the total 12 NILs from cross Pavon44:38/PBW550, almost all (besides NILs 13, 72, and 102) lines showed higher TGW than PBW550. The highest YD was observed for NIL102 (537.67g). For TGW, NIL88, 13, 97, and 11 showed lower values than the recurrent parent PBW550 among all the selected lines. For crossPavon44:38/DBW17, all the selected 9 NILs showed higher values for YD than the recurrent parent. 4 NILs showed lower values for the TGW than the recurrent parent DBW17. For TNpM, almost all (besidesNIL115, 128, and 108) NILs showed higher values than the recurrent parent DBW17.

There were 9NILs with the *Glu-B3+* gene being transferred by the donor parent Pavon40:9 in the background of PBW550 and DBW17 as the recurrent parents whose performance was superior/comparable to the respective recurrent parent (Table 3). In Pavon40:9XPBW550, NILs with *Glu-B3*+ gene, the YD value of 5 NILs (NIL-55, 6 7, 53, 56, and 64) was comparable to the recurrent parent. For TGW, NIL-56 and 67 showed higher values than the recurrent parent PBW550. Out of 6 NILs, 3 showed higher values (NIL-56, 64, and 67) for TNpM than the recurrent parent. In Pavon40:9XDBW17, NIL-58 and 61 showed higher value for YD than the recurrent parent. For TGW, NIL-61 and 57 values were high than the recurrent parent. The PH of all the selected NILs was in the moderate range of 75-110 cm. These selected NILs could be used to transfer *Glu-B3+* and *Sec-1-* genes into a single line to generate an improved version of the recombinant 1BL.1RS lines. 45 best performing BC_2_F_6_ NILs were selected and validated for the presence of glutenin/gliadin (42-50kDa) LMW proteins encoded by *Glu-B3* gene and and absence secalin protein (42-55kDA) on SDS base PAGE (Fig 2). 30 NILs were finally selected based on the yield related traits having 1RS chromosomal arm substituted with loci *Sec-1* and *Glu-B3* from wheat 1BS chromosomal arm.

## Discussion

The short arm of rye in 1BL.RS translocation has been widely used in bread wheat improvement programs worldwide for more than 50 years as a source of disease and pest resistance and higher grain yield potential (Oak and Tamhankar 2016). Besides the positive influence of this segment on yield and disease resistance, the 1RS translocation has been reported to be associated with severe defects in the bread-making quality such as poor mixing tolerance, dough stickiness, and low bread volume. This was mainly due to presence of *Sec-1* on its proximal end encoding tii-and γ-secalins proteins and absence of *Glu-B3/Gli-B1* loci encoding on its the distal end (Howell *et al.* 2014). Various methods of homeologous recombination by chromosome engineering and RNA interference for silencing the expression of secalins have been exploited to overcome such quality defects (Zhi et al. 2016). However, backcrossing seems to be simple and most effectively used method to replace two important loci on 1RS arm. In the current study, these two important loci on chromosome arm 1RS of elite wheat cultivars PBW550 and DBW17, were replaced with their wheat counterparts and NILs with improved 1RS arm and enhanced yield related traits were developed using combination of MAS and phenotypic evaluation focusing on yield and its contributing subcomponents.

The superiority of PBW 550 and DBW17 varieties in terms of quality parameters makes them the most suited as recipient for alteration in two targeted loci. PBW 550 is the medium short duration variety known for its bold grain and processing quality and was cultivated in almost all wheat growing regions of the country in the few years since its release in the year 2008 (Kaur et al, 2020). DBW 17, another full duration cultivar with appropriate plant height and lodging tolerance has protein content ranging between 11-12%. The high extraction rate in DBW17 flour to the tune of 70.4% enhances its industrial suitability for more flour recovery (Singh et al. 2007).

All the markers used in current study had been reported to have strong linkage with the corresponding loci making the making the MAS most effective as reported by Sharma et al. 2018. Moreover, for background recovery of recurrent parent as such and 1RS/1BL chromosome, a comparatively higher number of markers were deployed from 1BL chromosomal arm as more recombinants for this chromosome were expected to be selected while doing MAS for targeted loci. Two backcrosses successively recovered the recurrent genotypes of PBW550 and DBW17.

Based on phenotypic selections across the two-year trials, 30 NILs with superior performance for yield-related traits than their respective recurrent parents were selected (Table 3). Despite large GxE interaction of the lines across the years (Fig. 4), many genotypes showed better performance than the recurrent parents across both years (Table 3). Large genetic variability among the NILs was detected within each of the two environments, which aided in the selection of better-performing lines (Singh et al. 2018; Dabi et al. 2019).

The exploitation of variability in the form of Phenotypic Coefficient of Variation (PCV), Genotypic Coefficient of Variation (GCV), and heritability is essential for the proper selection of lines for any breeding program (Neelima et al. 2020). Moderate to high GCV and PCV for PH, TNpM, TGW, HI, and YD indicated that the phenotypic selection could improve these traits. Osman et al. (2012) reported that the environmental effect on any trait is indicated by the magnitude of the differences between the genotypic and phenotypic coefficients of variation; large differences reflect a large environmental effect, whereas small differences reveal a high genetic influence. In this study, the small differences between the PCV and GCV for most of the traits, *i.e.,PH*, TNpM, TGW, YD, and HI, represented only a small degree of environmental influence phenotypic expression of these characters in the respective years. It also suggests that selection based on these characters would be effective for future crossing programs. The heterogeneity coefficients (GCV, PCV) values alone are insufficient to determine the heritable portion of variance passed on from generation to generation, as expressed by broad-sense heritability (Lush 1949). Also, the broad-sense heritability estimates the quality of the breeding program for the traits and the environments. In the present study, moderate to high broad-sense heritability of the studied traits indicated a higher contribution of genotypic component of variation for the traits than the environmental effect across the various blocks in each year trial (Table 3). The estimate of heritability is a predictive measure scrutinizing the reliability of the phenotypic data, and genetic advance as percent of mean is evidence of the expected result of the application of a selection pressure on the pertinent population. Hence, the heritability along with GAM offers a more consistent index for the selection value (Hanafi et al.2020). The GAM of second year was found to be higher than the first which indicates positive effect of the selection leading to selection of superior lines.

Overall, the TNpM and TGW had a positive correlation with the YD. TGW is directly related to the grain yield and milling quality of the grain and impacts the seedling vigor and growth, indirectly affecting the yield (Botwright et al. 2002; Feng et al. 2009; Wu et al. 2018). Higher grain weights are positively associated with longer grain filling duration attributed to timely flowering and high grain filling volume (Zhang et al. 2010; Okami et al. 2016). As identified in the present study, this positive correlation has been reported in various studies (Kumar et al. 2014; Bhutto et al. 2016; Birhanu et al. 2017; Reddy et al. 2021). TNpM was found to have a positive correlation with yield across the years. Bhutto et al. (2016) published similar findings, demonstrating that an increase in the number of tillers leads to a proportionate increase in yield per plot. HI had shown positive correlation with YD as previously documented in different studies (Yang and Zhang 2010; Duan et al. 2018; Ziang et al. 2019). The positive association between yield and HI could be of major significance in encouraging breeders in their exploration for increased yield in wheat varieties (Foulkes et al. 2009; Aranjuelo et al. 2013; Duan et al. 2018).

The nature of gene action and the number of genes controlling the trait is usually measured by the critical analysis of distribution properties by third order statistics such as skewness and kurtosis which are more important than the first and second order statistics that unravel only the interaction effects (Rani *et al.* 2016). Skewness indicates the cluster of deviation above and below the value of central tendency and defines the extent of deviation in the distribution of trait values and thus could aid in detection of varying effects like additive effects, dominance, and also epistasis. Positive skewness would indicate the traits to be controlled by dominant and complementary gene action whereas a negative skewness would indicate the traits to be controlled by dominant and duplicate epistasis (Neelima et al. 2020). All the traits exhibited positive skewness for both years suggesting that the genetic gain obtained is through intense selection (Table 1).

Kurtosis indicates the level of peakness over the population with a leptokurtic distribution would mean that the trait in question is controlled by fewer genes whereas a platykurtic distribution would mean that the trait is governed by many genes (Savitha and Kumari 2015). The decrease in leptokurtic value of PH for the second year suggests that the selection led to removal of introgression lines with outlier trait values. The leptokurtic values for second year could also be because of the decrease in the number of the lines that resulted in overall less distribution.

Principal component analysis is an exploratory tool for data analysis. It offers details about traits by elucidating the population’s maximum variability in the given environments. High GxE effect was observed across the years depicted by lower variability (51.3%) explained by the eigen vectors in the first two principal components (Fig. 4). Similar results have previously been reported where lower variability explained by first two components has been associated to large number of genic interactions among the traits which is further complexed by the additive effect of the genes involved for each trait (Hailegiorgis et al.. 2011; Degewione and Alamerew 2013; Nielsen et al.. 2014; Mohibullah et al.. 2017; H. Wani et al.. 2018; Kiran et al.. 2021). TGW was shown to be highly dependent on HI and TNpM and least dependent on SN and SL in the first two PCAs. Similarly, SN was heavily reliant on SL. On the other hand, YD was based on all of the characters analysed, with HI, TGW, and PH being the most dependent variables (Tshikunde et al. 2019). Since the PCA revealed that YD was influenced by all of the characters examined in this experiment to varying degrees, the SEM revealed that the main direct contributors to YD were the TNpM, HI and TGW (Fig. 6). TGW also have an indirect effect on YD through HI. The PH has indirect effect on YD through HI and TNpM (Bhutta et al. 2006). SL has a direct negative effect on YD and SN and directly affects TGW and indirect negative effect on YD, as explained in PCA (Iftikhar et al. 2012).

Agronomic evaluation of the NILs identified 30 lines that demonstrated substantial improvements for TGW, YD and TNpM. Highest number of NILs (11) with*Sec-1-* gene (Pavon44:38XPBW550) having a high TGW, TNpM and many of these had high YD. Ten lines of Pavon44:38XDBW17 were chosen for their high YD. Similarly, nine NILs *withGlu-B3+* gene (Pavon40:9XPBW550/DBW17) showed better performance for YD, TGW and TNpM. Two lines inDBW17 background had higher YD. Of seven lines in PBW550 background, two have high TGW and the three have high TNpM and YD. The average YD in the second year was less than that of the average YD for the first year. This could be because of the environment effects which have not been calculated in current study. Over all the yield performance of the lines with *Sec-1-/Glu-B3-* was better than those with *Sec-1+/Glu-B3+* indicating the secalin locus is not contributing towards yield and its removal was rewarding also (Kaur et al. 2017).

The marker-based selection of the improved recombinant lines with *Glu-B3*+and/or *Sec-1-* could be evaluated for quality-related traits of sedimentation value (cc), gluten index, dry gluten (%), hectolitre weight (kg/hl), protein content (%), and grain hardness (kg). Concerning farmers’ perspective, these lines are already thriving in providing increased yield as they are resistant to various rust gene(s) and have good root architecture gene(s). Further, for consumer usage, quality is a primary concern these days in response to change in living standards, and improved nutritional values of these lines could be a boon at the commercial level.

## Acknowledgements

The financial support provided by the Department of Biotechnology, Ministry of Science and Technology, Government of India in the form of grant No. BT/PR10886/AGII/106/934/2014 is gratefully acknowledged.

## Author’s Contributions

RK, PK and DK developed the material, conducted marker assisted selection. RK, SK and RS done background selection. GSD and AKaur done phenotypic analysis; AKumar helped in validation on SDS-PAGE. GSV and SKG helped in development of material and phenotypic evaluation; SK* and Kaur supervised the study and helped in writing the manuscript; SK* and PC designed the study, provided the basic genetic material, and finalized the manuscript. All the authors have read the manuscript and approved it.

